# Dynamic changes in clonal architecture during disease progression in follicular lymphoma

**DOI:** 10.1101/181792

**Authors:** Christoffer Flensburg, Tobias Sargeant, Astrid Bosma, Roelof J. C. Kluin, Robby E. Kibbelaar, Mels Hoogendoorn, Warren S. Alexander, Andrew W. Roberts, René Bernards, Daphne de Jong, Ian J. Majewski

## Abstract

Follicular lymphoma (FL) is typically a slow growing cancer that can be effectively treated. Some patients undergo transformation to diffuse large B cell lymphoma (DLBCL), which is frequently resistant to chemotherapy and is generally fatal. Targeted sequencing of DNA and RNA was applied to identify mutations and transcriptional changes that accompanied transformation in a cohort of 16 patients, including 14 with paired samples. In most cases we found mutations that were specific to the FL clone dominant at diagnosis, supporting the view that DLBCL does not develop directly from FL, but from an ancestral progenitor. We identified frequent mutations in *TP53*, cell cycle regulators (cyclins and cyclin-dependent kinases) and the PI3K pathway, as well as recurrent somatic copy number variants (SCNVs) on chromosome 3, 7 and 17p associated with transformation. An integrated analysis of RNA and DNA identified allele specific expression changes in oncogenes, including *MYC*, that could be attributed to structural rearrangements. By focusing on serial samples taken from two patients, we identified evidence of convergent tumour evolution, where clonal expansion was repeatedly associated with mutations targeting the same genes or pathways. Analysis of serial samples is a powerful way to identify core dependencies that support lymphoma growth.

## INTRODUCTION

Follicular lymphoma (FL) arises from the germinal centre B cell compartment and represents the most common form of non-Hodgkin’s lymphoma. The chromosomal translocation t(14;18), which drives high level expression of the pro-survival molecule BCL2, is a hallmark of FL and is present in more than 90% of cases (1-3). Although the translocation provides a strong survival stimulus, it is not enough to cause the disease, indicating that additional cooperating events are required (4, 5).

FL is generally an indolent disease, but around 30% of patients will subsequently develop an aggressive transformed lymphoma, typically Diffuse Large B cell Lymphoma (DLBCL) (6). With modern management, transformation is the most common cause of lymphoma-related death, and so detailed understanding of its pathobiology is needed. A range of techniques have been applied in the search for genetic events that initiate transformation, including karyotyping (7), structural analysis with single nucleotide polymorphism (SNP) arrays (8), monitoring patterns of somatic hypermutation and immunoglobulin rearrangement (9, 10), and, more recently, genomic profiling (11-15). These studies reveal a diverse array of mutations that contribute to disease progression, but also demonstrate that the clonal architecture of FL is highly complex. Rather than a direct linear progression from FL to DLBCL, transformation typically occurs via a more ancestral progenitor.

In this study we examine genetic and transcriptomic changes in FL patients at multiple points during their disease course. Our analysis tracked somatic single nucleotide variants and small insertions and deletions (collectively referred to here as somatic single nucleotide variants, SSNVs), and somatic copy number variants (SCNVs), to extract a detailed view of the clonal architecture of the disease. We identified mutations associated with progression from FL to DLBCL that affect key regulators of the cell cycle and the PI3K pathway. By studying multiple relapses from an individual patient, we saw evidence of convergent disease evolution, with the same genetic alteration occurring independently, multiple times, and the repeated acquisition of distinct mutations that activate the same cellular pathways.

## METHODS

### Patient selection

The clinical series includes 16 patients, including some that were previously profiled in a large gene expression study (16), who presented with FL and subsequently developed transformed disease (interval 2-184 months, median 36 months) and for whom cryopreserved biopsy samples were available **(Supplementary Table 1)**. Biopsy samples of both disease phases were available for 14 patients, with serial samples available for 2 of these. All patients were treated at the Netherlands Cancer Institute and at the Friesland Medical Centre between 1993 and 2009. All protocols for obtaining and studying human archival tissues and patient data were approved within the local ethical procedures at the Netherlands Cancer Institute [METC16.1404] in compliance with the Code for Proper Secondary Use of Human Tissue in the Netherlands (17).

DNA and RNA were extracted from cryopreserved tumour tissue. DNA was extracted using the DNEasy Blood & Tissue kit according to the manufacturer’s instructions (Qiagen, Hilden, Germany). Total DNA concentration was assessed using absorbance at 260 nm on spectrophotometer. Total RNA was extracted using TriZol according to the manufacturer’s instructions (Life Technologies, Carlsbad, CA, USA). Quality assessment for RNA was performed with a Bioanalyzer (Agilent, Santa Clara, CA, USA).

### Capture enrichment and sequencing

Genomic DNA was used to sequence a set of 630 genes, enriched for kinases and common cancer genes **(Supplementary Table 2)**, for all biopsy samples from all patients. Indexed genomic DNA fragment libraries were prepared using the TruSeq genomic DNA library kit (Illumina, San Diego, CA, USA). When RNA was available (11 patients) capture enrichment was also performed with RNA sequencing (RNAseq) libraries. Indexed RNA sequencing (RNAseq) libraries were constructed from poly-A enriched RNA using the TruSeq mRNA library preparation kit (Illumina). DNA and RNA sequencing libraries were enriched through capture with the human kinome DNA capture baits (Agilent), as described previously (18). Six libraries were pooled together for each capture reaction. After capture the enriched libraries were sequenced on an Illumina HiSeq2000, using a paired end 55bp protocol. Sequencing data has been deposited in the European Genome-phenome Archive [EGAS00001002175].

### Sequence alignment, variant calling and copy number profiling

DNA reads were aligned to hg19 using bwa mem v0.7.10-r789 (19) with default settings, and RNA reads were aligned with Tophat (20) v2.0.12. Variant calling for DNA was performed with SAMtools version 0.1.19-44428cd followed by Varscan (21) v2.3.6. Detailed analysis settings are available in supplementary methods. Liberal settings were used for variant calling and the results were then filtered to remove low quality variants and mapping artefacts.

Variant analysis and copy number calling was performed with superFreq (22). A more detailed description of the analysis pipeline is available in supplementary methods. First, candidate variants were classified as either germline SNPs, SSNVs, or false positives, which involved assessing sequencing and alignment quality scores, and by comparison to dbSNP (23) and a reference set of 17 normal samples. We also used a Dutch population study to identify additional germline SNPs (24). SCNVs were identified using changes in allele frequencies at heterozygous germline SNPs and changes in coverage at capture regions, relative to a pool of normal control samples. SSNV and SCNV clonalities were monitored in serial samples to track clonal selection during disease progression. All variants classified as SSNVs were annotated using Ensembl Variant Effect Predictor version 75 (25). We classified mutations as being present in FL, DLBCL, or both.

### Clonality assessment for patients with multiple samples

The clonal analysis of patient 4 and 14 was made with output from superFreq, that combines and tracks SSNVs and SCNVs across samples. For each mutation both the clonality and uncertainty were assessed across all samples from that patient. Local copy number was used to inform clonality estimates for SSNVs. The resulting clonality estimates were clustered to group mutations that show similar patterns of prevalence, enabling the identification of events that occur in the same cell population. The code for the analysis and figure generation is available in the supplementary materials.

### Differential expression and gene set enrichment analysis (GSEA)

RNAseq libraries were also captured with baits targeting the kinome. We used featureCounts (26) together with limma (27) and voom (28) to perform the differential expression analysis. The XRank algorithm (29) was used to calculate a corrected log fold change (LFC) that was used as a ranking statistic for GSEA (30). When assessing expression changes in individual samples we compared expression in that sample to all FLs (when the sample was a FL, it was compared to all other FLs) using the same methods.

### Identification of shifted allele ratios in RNA

Allele frequencies were assessed at heterozygous germline SNPs and compared between DNA and RNA to identify significant differences using Fisher’s exact test. Significance was Benjamini-Hochberg corrected for multiple testing based on the number of heterozygous germline SNPs. We concentrated on genes that included at least one SNP where the non-reference allele frequency increased in the RNA. This requirement helped to limit the influence of reference mapping bias around splice sites, but does limit sensitivity. Genes identified in this analysis were manually inspected in IGV to ensure that the signal was not due to alignment or quality issues.

### Immuno-histochemistry and fluorescence in situ hybridisation (FISH)

Formalin-fixed paraffin embedded (FFPE) tissue sections of representative biopsy samples were immunostained according to standard procedures, including antigen retrieval (CC1) on Ventana Benchmark Ultra (Ventana, Tucson, USA) and using antibodies against MYC (EP121, Epitomics, Burlingame, CA) and TP53 (DO-7, DAKO, Glostrup, Denmark). Immunohistochemical results were visually assessed and quantified in 10% increments. Interphase FISH was performed on tissue sections of the same FFPE biopsy samples using break-apart probes (Vysis, LSI MYC break apart, Des Plaines, IL) and evaluated under a fluorescence microscope (Zeiss, Göttingen, Germany). All assessments were performed using standard H&E and immunohistochemistry for diagnostic purposes as a reference.

## RESULTS

### Integrated mutational profiling during disease progression in follicular lymphoma

We sought to identify mutational events that occurred early in the disease, that are present in all cancer cells, and disease phase-specific events. We assessed the number of mutations in each sample and used paired samples to classify mutations as early events, or disease phase-specific events **(Figure 1, Supplementary Table 3)**. On average 16 mutations were identified in each FL at diagnosis. For 11 patients we observed mutations that were unique to the FL, indicating that the DLBCL did not emerge directly from the FL clone that was dominant at diagnosis, but from an earlier common progenitor. In the remaining 3 patients no mutations were detected that were unique to FL, allowing for the possibility of direct progression from FL to DLBCL. As anticipated, we observed more phase-specific mutations in DLBCL than in FL (on average 10 compared to 2); these events are of special interest because they may identify mechanisms that contribute to transformation.

**Figure 1:**
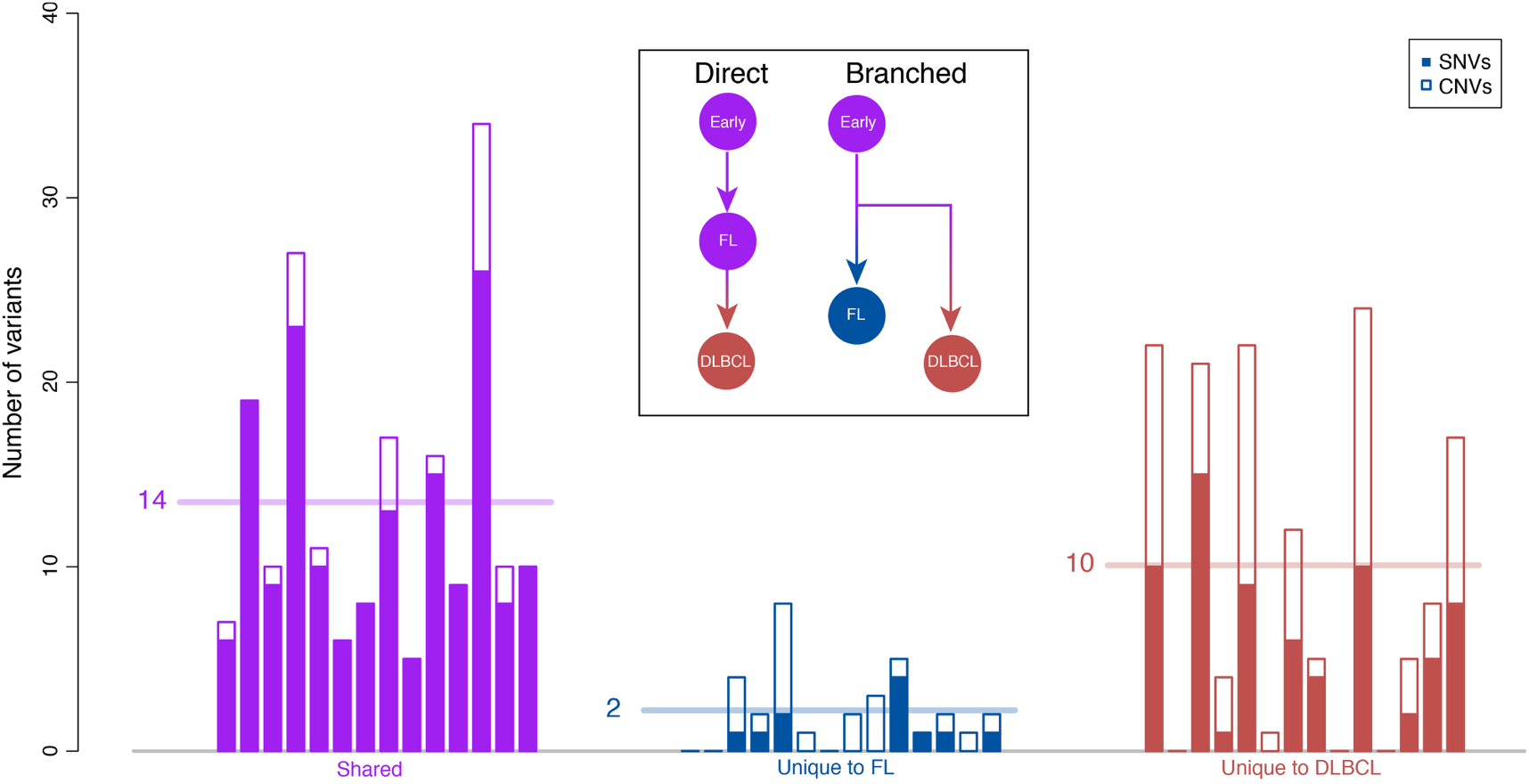
Assessment of mutation load in FL and DLBCL. Patients 2-15 are shown in order from left to right. The number of mutations detected in each sample is plotted and the average number is indicated by a horizontal line. SSNVs were included if they had an allele frequency >20%, and were classified as unique to FL or DLBCL if the allele frequency in the other disease phase was both <20% and significantly different (Benjamini Hochberg adjusted p-values from Fisher’s exact test below 5%). SCNVs were included if they spanned at least 10 Mbp and had a clonality greater than 40%. SCNVs were classified as unique if they did not overlap a SCNV with similar coverage and genotype in the other disease phase. Inset, the presence of unique mutations within the FL supports a branched model where the DLBCL originates from an earlier progenitor population.

Analysis of genome-wide copy number profiles identified recurrent SCNVs associated with transformation to DLBCL, including gain of chromosome 3 and 7, and loss of 6q, 9p and 17p **(Figure 2A)**. Additionally, biallelic mutations that combine SSNVs and SCNVs were identified in *TP53*, *CDKN2A*, *PTEN* and *DGKZ* **(Supplementary Figure 1)**. Three patients acquired missense or nonsense mutations in *TP53* (Patient 2: C277F, Patient 6: R342X and Patient 15: S260C) during transformation to DLBCL that were concurrent with loss of the other allele, while a fourth patient exhibited loss of one allele. Immunohistochemistry was used to confirm stabilisation of TP53 in two cases with mutations

**Figure 2:**
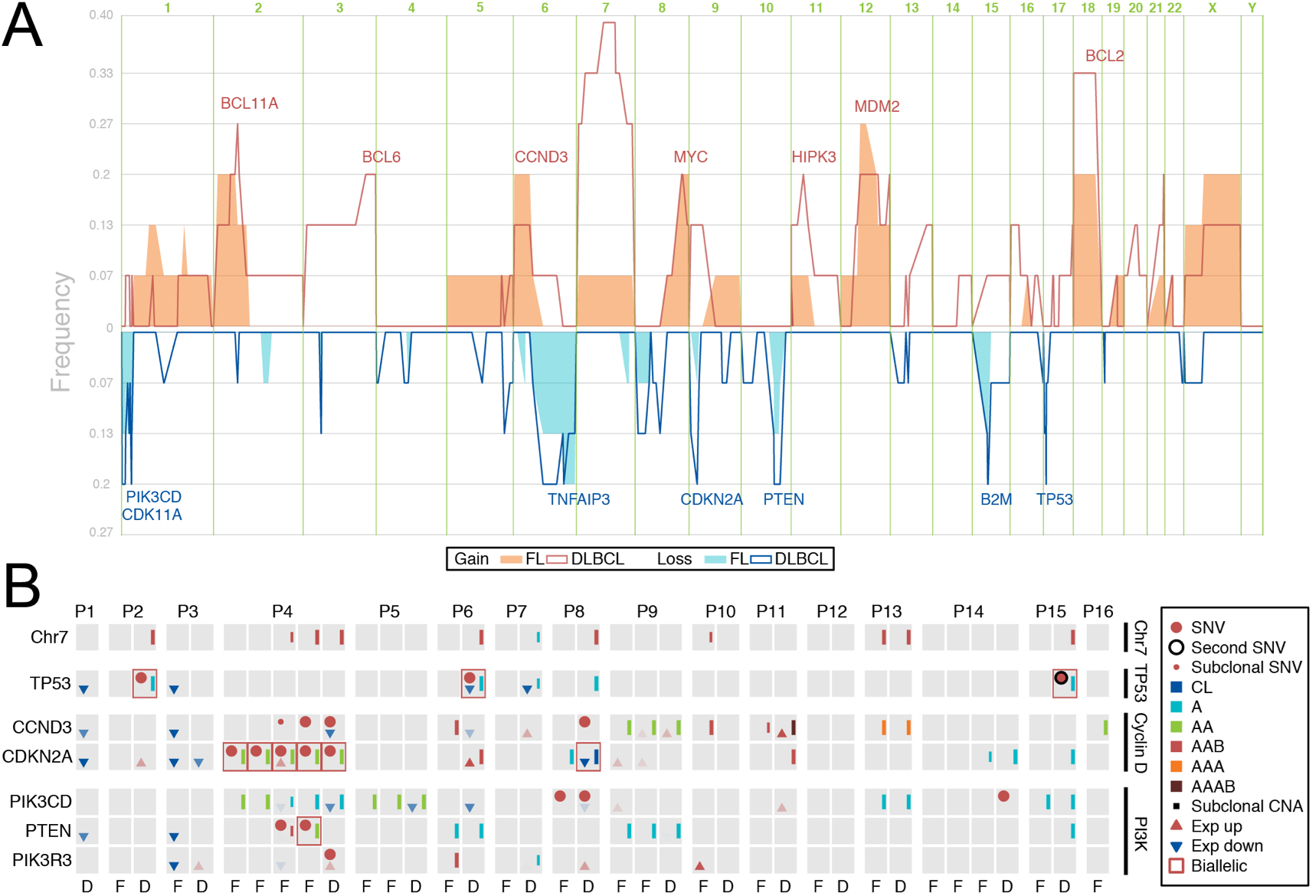
Copy number and mutation profiling for key pathways. A. Copy number alteration frequency in FL and DLBCL. Gains (in red, at top) and losses (in blue, at bottom) are presented as frequencies, assessed separately for FL and DLBCL. Some genes implicated in FL/DLBCL biology have been labelled. B. Mutations and differential expression were assessed for genes in three key pathways: TP53, Cyclin D (*CCND3*, *CDKN2A*) and mTOR (*PIK3CD*, *PTEN*, *PIK3R3*). Each box represents an individual sample, displayed chronologically. Detection of a non-synonymous SSNV is indicated at the top left of each box. SCNVs are represented by coloured lines on the right side, which indicate copy number and genotype (CL: complete loss, A: loss of one allele, AA: CNN-LOH, AAB: gain of one allele, AAAB: gain of two copies of one allele, AAA: three copies of the same allele). Changes in expression were assessed relative to all other FL samples and are displayed in the bottom left corner of each box (arrows indicate direction, transparent arrows represent LFC >0.5, opaque arrows represent LFC ≥1). RNAseq was un available for P12 to P16.

**(Supplementary Figure 2)**. Biallelic mutations in *CDKN2A* were detected in two patients. While mutations in *CDKN2A* were typically acquired upon transformation (Patient 8, Patient 14 and Patient 15), inactivation of *CDKN2A* was an early event in Patient 4, as all samples from that patient shared the same nonsense mutation (CDKN2A Y129X) that became homozygous due to copy number-neutral LOH (CNN-LOH). Both patients that had complete inactivation of *CDKN2A* also carried stabilising mutations in *CCND3* (Patient 4: L292R and Patient 8: Q260X), consistent with previous work suggesting synergy between these mutations(31). One patient had biallelic inactivation of *PTEN* through a homozygous missense mutation and CNN-LOH (Patient 4 FL 2006: V166I), while three additional patients displayed loss of one allele. The mutation status and expression of *TP53*, *CDKN2A*, *PTEN* and other pathway components was assessed across the entire cohort **(Figure 2B)**.

**Supplementary Figure 1:**
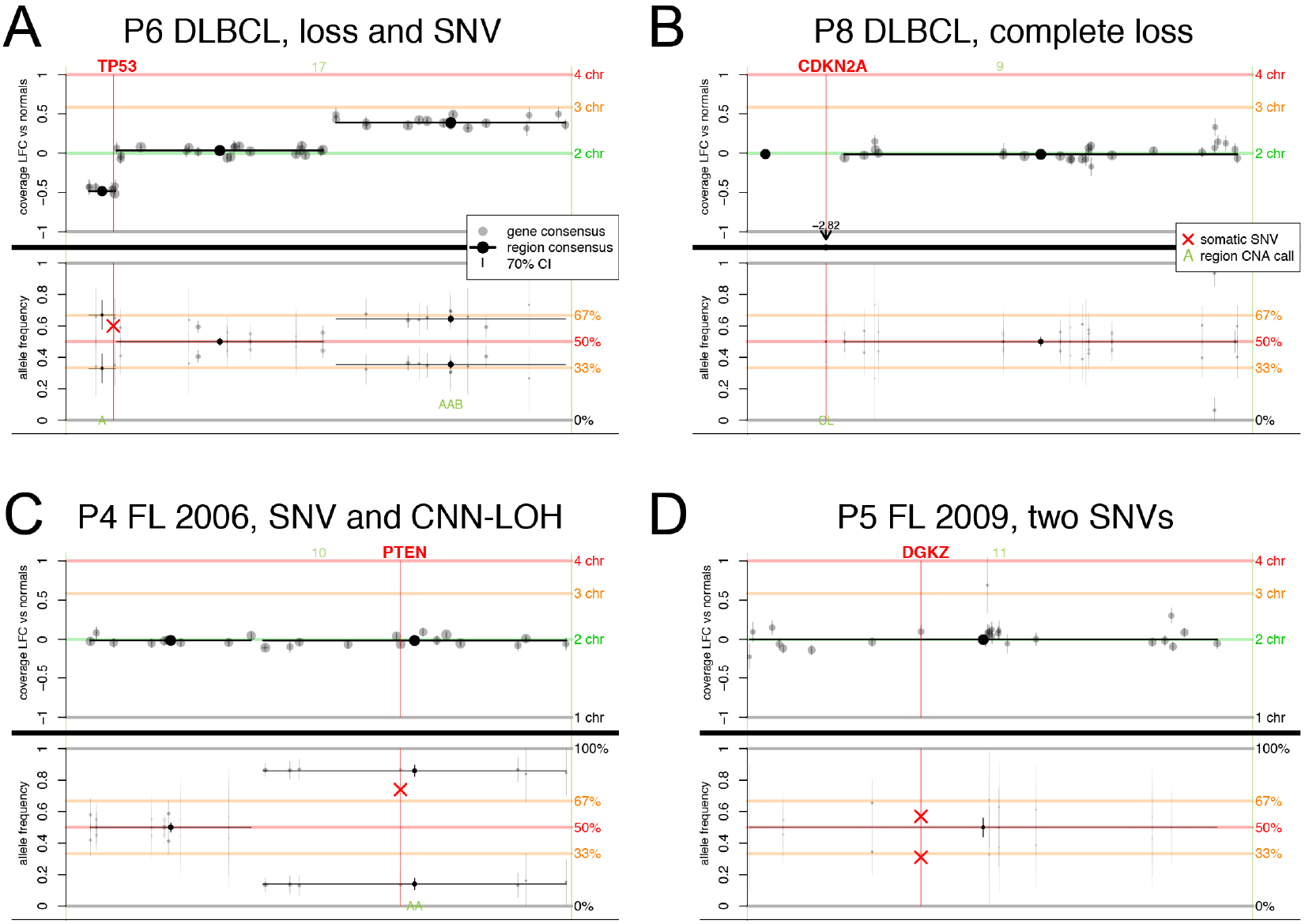
Biallelic mutations in key cancer genes. Examples of biallelic mutations in (a) *TP53*: R342X and loss in P6 DLBCL, (b) *CDKN2A*:, complete loss in P8 DLBCL, (c) *PTEN*: V166I which becomes homozygous due to CNN-LOH in P4 FL 2006 and (d) *DGKZ*: E607K and a frameshift (+C) in P5 FL 2009. The top panel of each figure shows the LFC of the coverage compared to the pool of normal samples. In panel (b) the LFC of CDNK2A is outside the scale so the value is presented. The lower panel shows the allele frequency for SNPs used in the copy number determination and for non-synonymous SSNVs detected in the genes of interest, with the later represented with red crosses. Genotype calls are provided for each clustered region in green font (CL: complete loss, A: loss of a single allele, AAB: gain of one allele, AA: CNN-LOH). Individual genes are represented with points and horizontal lines are used to denote SCNV regions, that is, groups of genes with the same copy number status. Error bars represent 70% confidence interval (Supplementary Methods).

**Supplementary Figure 2:**
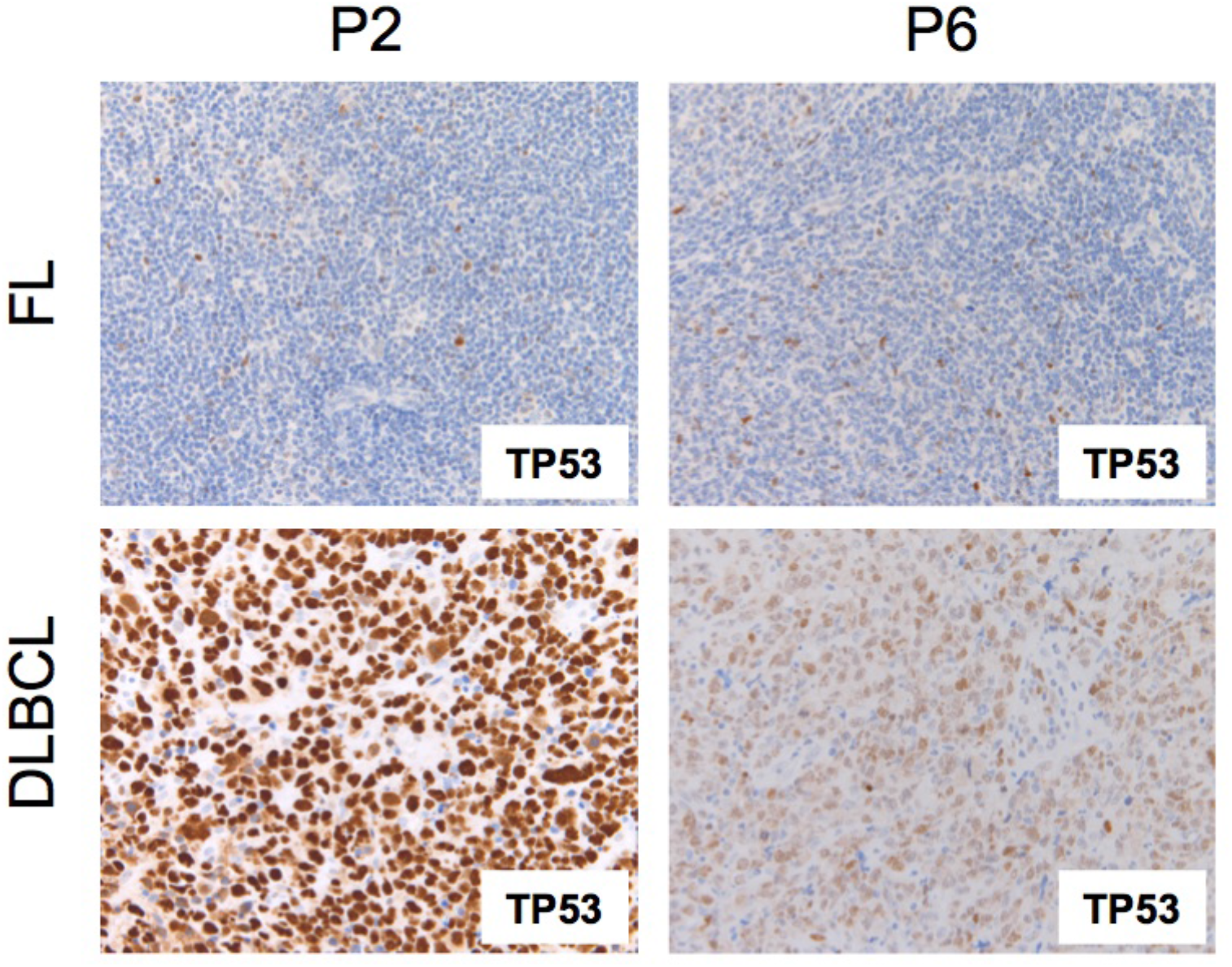
Elevated TP53 protein in DLBCL cases that carry mutations. TP53 levels were assessed in matched FL and DLBCL samples from Patient 2 (P2) and Patient 6 (P6) using immunohistochemistry. The proportion of cells that stained positively was markedly elevated in the DLBCL stage, consistent with the acquisition of loss of function mutations in TP53.

*Serial sampling identifies dynamic changes in clonal architecture during disease progression.*

For Patient 4 and Patient 14 we were able to assess samples taken over the course of treatment, which allowed for a deeper understanding of the relationship between each relapse. We calculated the clonality – the proportion of cells in each sample that carry the mutation – and grouped mutations that showed similar prevalence over time. These groups can be used to infer the evolutionary history of the cancer.

For Patient 4 we had access to diagnostic material (2003 FL) and three subsequent relapses (2005 FL, 2006 FL and 2008 DLBCL), and could identify seven groups of mutations with different prevalence. We were able to generate a tree to represent the evolutionary history of the disease based on the mutation groups **(Figure 3)**. This tree was supported by mutations in the 5’UTR of BCL2 resulting from somatic hypermutation that were detected in matching RNAseq **(Supplementary Figure 3)**. We identified two distinct mutations in *CDKN2A* that were present clonally at all time points (black line in Figure 3), indicating that the mutations occurred early in the history of the cancer. The diagnostic FL sample also exhibited CNN-LOH over a 12Mbp region on chromosome 1p, which was present at lower levels in 2005 and lost at later time points, indicating extinction of the FL clone dominant at diagnosis (dark blue line in Figure 3). All relapse samples shared a missense mutation in CCND3 (L292R) that has been shown to stabilise the protein (31). The *CCND3* mutation appeared in 2005 FL at 40% clonality, but became the dominant population in 2006 and 2008. There are eight mutations (SSNVs and SCNVs) that show the same pattern, suggesting each relapse originated from a common progenitor that was distinct from the clone that gave rise to the FL at diagnosis.

**Figure 3:**
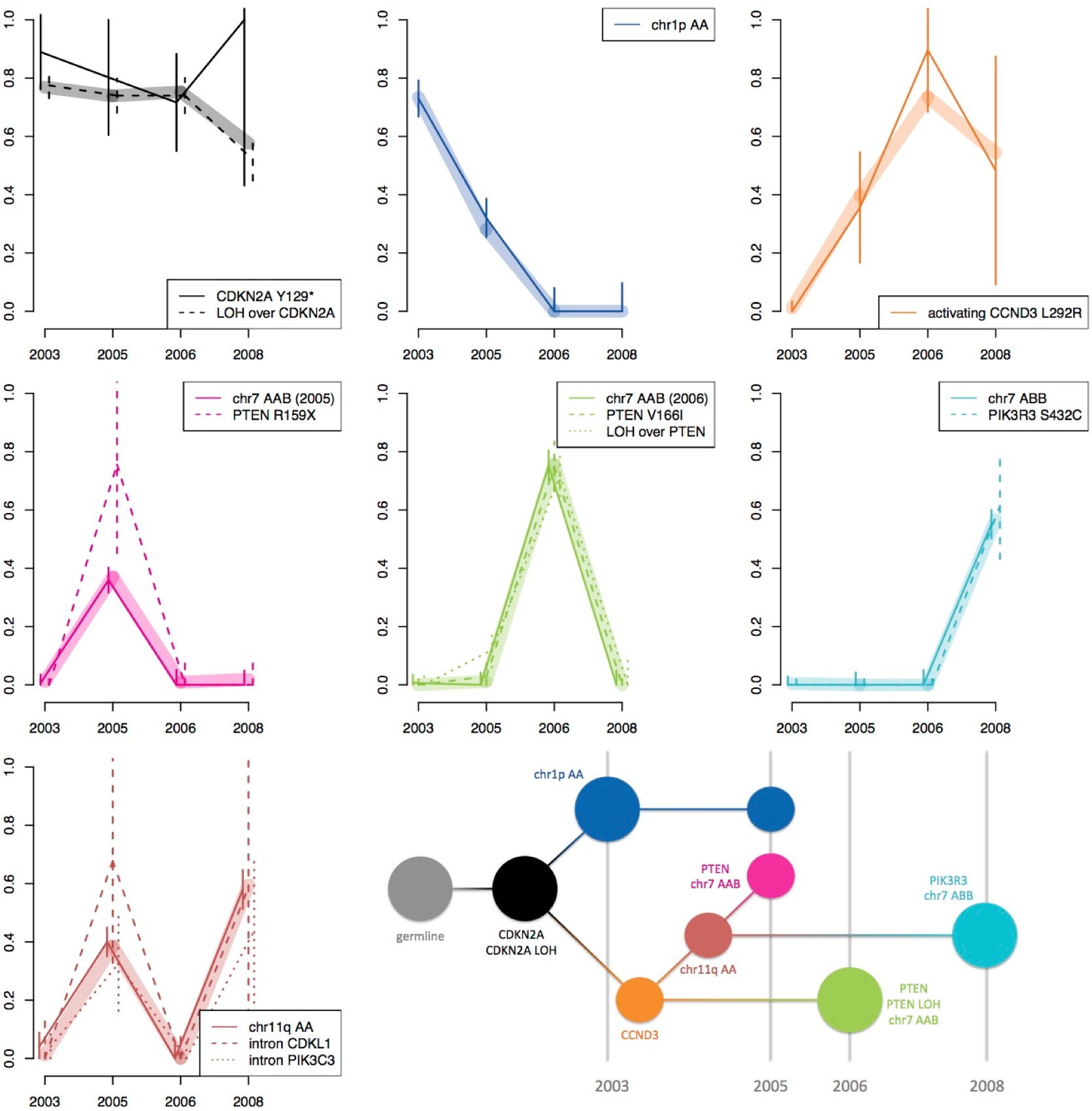
Dynamic changes in clonal architecture during disease progression in a single patient. Mutations with similar patterns of prevalence were grouped. The clonality of each mutation group is plotted for diagnosis and relapse samples: thick lines show the consensus clonality (weighted mean of the grouped mutations), thin lines highlight individual mutations in the clone and error bars represent a 70% confidence interval. The pattern of mutations was used to infer the evolutionary history of the cancer. These relationships are represented in a tree structure and some key mutational events are highlighted. The cyan clone gained a different allele of chromosome 7 than the pink and green clones, which is shown as “chr7 ABB” as opposed to “chr7 AAB”.

**Supplementary Figure 3:**
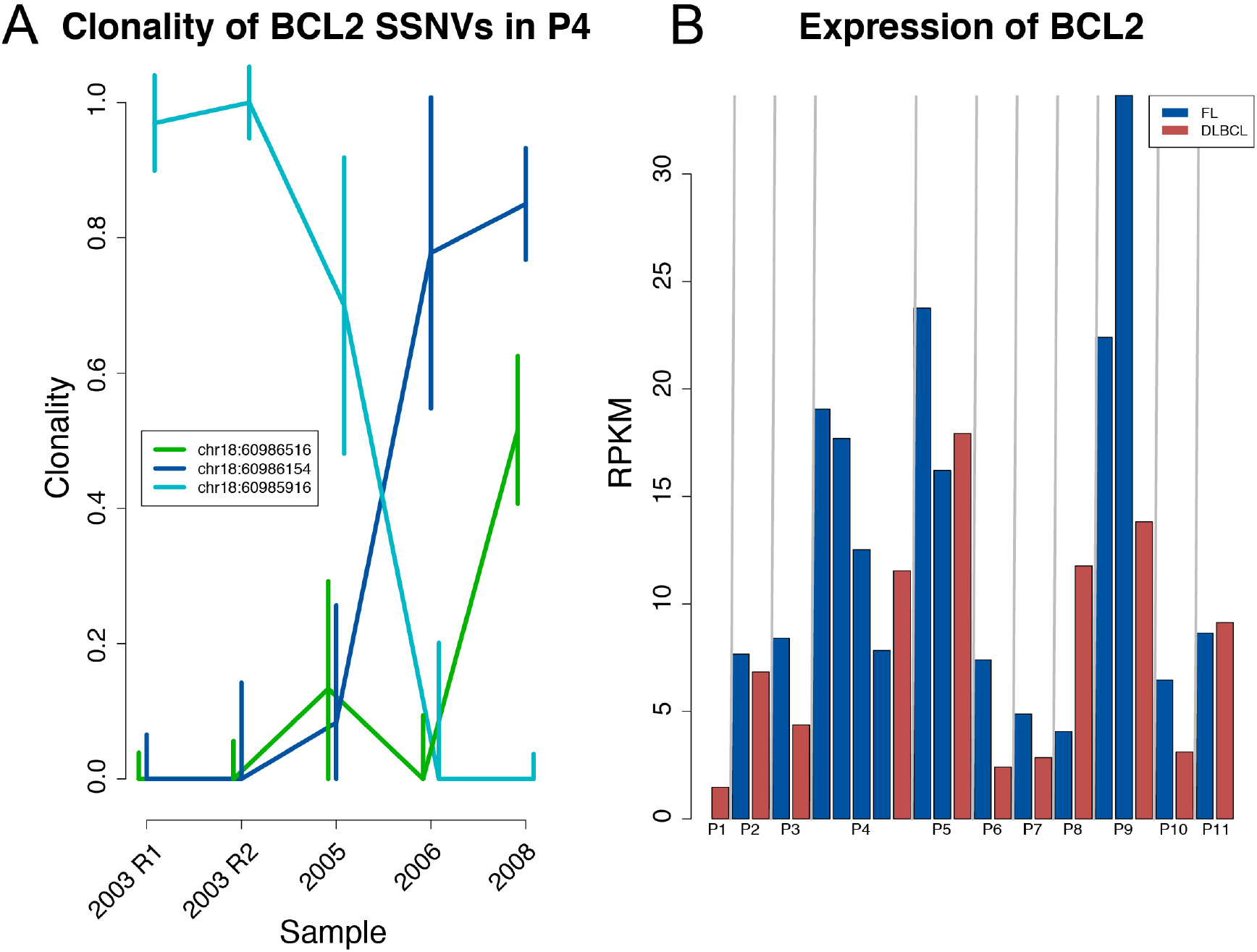
Clonality of three mutations in BCL2 resulting from somatic hypermutation. A. The clonality of three SSNVs detected in *BCL2* in five RNA samples from Patient 4. Subpopulations within a sample may express *BCL2* at different levels, which impacts on the clonality estimate. For example, the clonality estimate of the BCL2 SSNV present at diagnosis (cyan) is 70% in 2005 in the RNA, but is only observed at 30% clonality in the matching DNA (blue in Figure 5). This disparity results from *BCL2* being more highly expressed in the diagnostic FL than in the relapse. New SSNVs detected in 2005 become dominant at later time points, consistent with the pattern observed with DNA. B. The expression level of *BCL2* as RPKM across the samples.

When we examined the mutations that were specific to each relapse from Patient 4 we identified multiple mutations targeting the PI3K pathway. The first relapse in 2005 carried a nonsense mutation in *PTEN* (R159X), the second relapse in 2006 acquired biallelic inactivation of *PTEN* (V166I, which has become homozygous due to CNN-LOH) and the third relapse had a missense mutation in the regulatory component *PIK3R3* (S432C) **(Figure 3)**. All relapse populations had separately gained chromosome 7, as inferred from differences in the genotype (gain of either the maternal or paternal chromosome 7) and the clonal architecture **(Supplementary Figure 4 and Supplementary Methods)**. This suggests that gain of chromosome 7 and activation of the PI3K pathway cooperate to mediate rapid outgrowth on a background of biallelic loss of *CDKN2A* and stabilised *CCND3*. Comparison between the relapse FL sample and the DLBCL from Patient 14 demonstrated each had lost a single allele of *CDKN2A*, but these were distinct events **(Supplementary Figure 5)**. Collectively these findings suggest convergent evolution within the disease, identifying common pathways that support clonal expansion.

**Figure 4:**
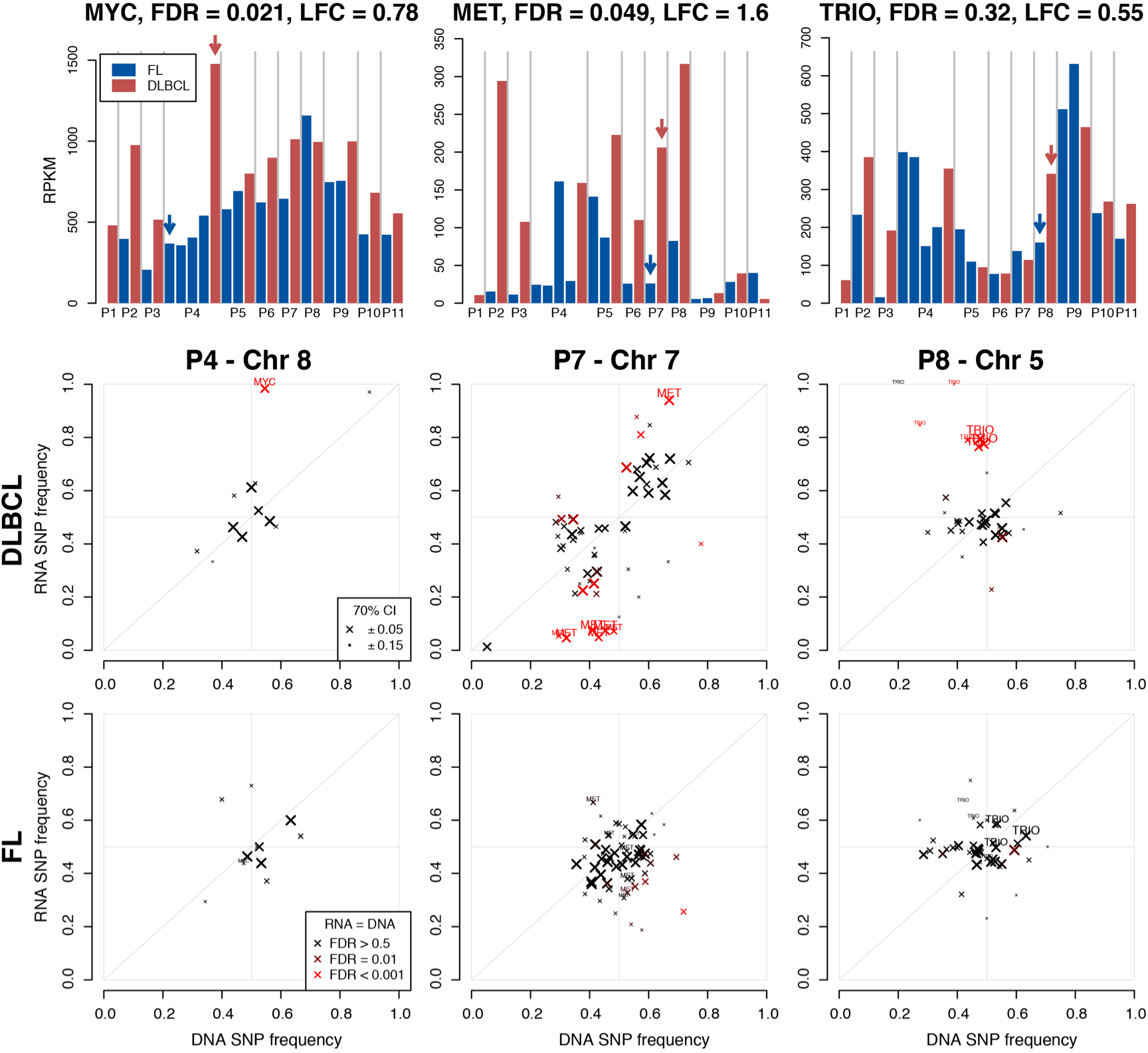
Shifted allele frequencies in RNA for *MYC*, *MET* and *TRIO*. Top panels: The expression of *MYC*, *MET* and *TRIO* is expressed as reads per kilobase per million (RPKM) for each sample. FDR and LFC refer to the differential expression between DLBCL and FL. Three patients show evidence of shifted heterozygous allele ratios in the transformed samples. Middle panels: The allele frequency is shown for heterozygous germline SNPs in DNA and matched RNA for DLBCL samples. The accuracy of the allele frequency, influenced by coverage, is represented by point size (see supplementary methods). SNPs with significantly different frequencies between DNA and RNA are coloured red and those from the three genes of interest are labelled. Corresponding plots for FL samples are shown in the bottom panels.

**Figure 5:**
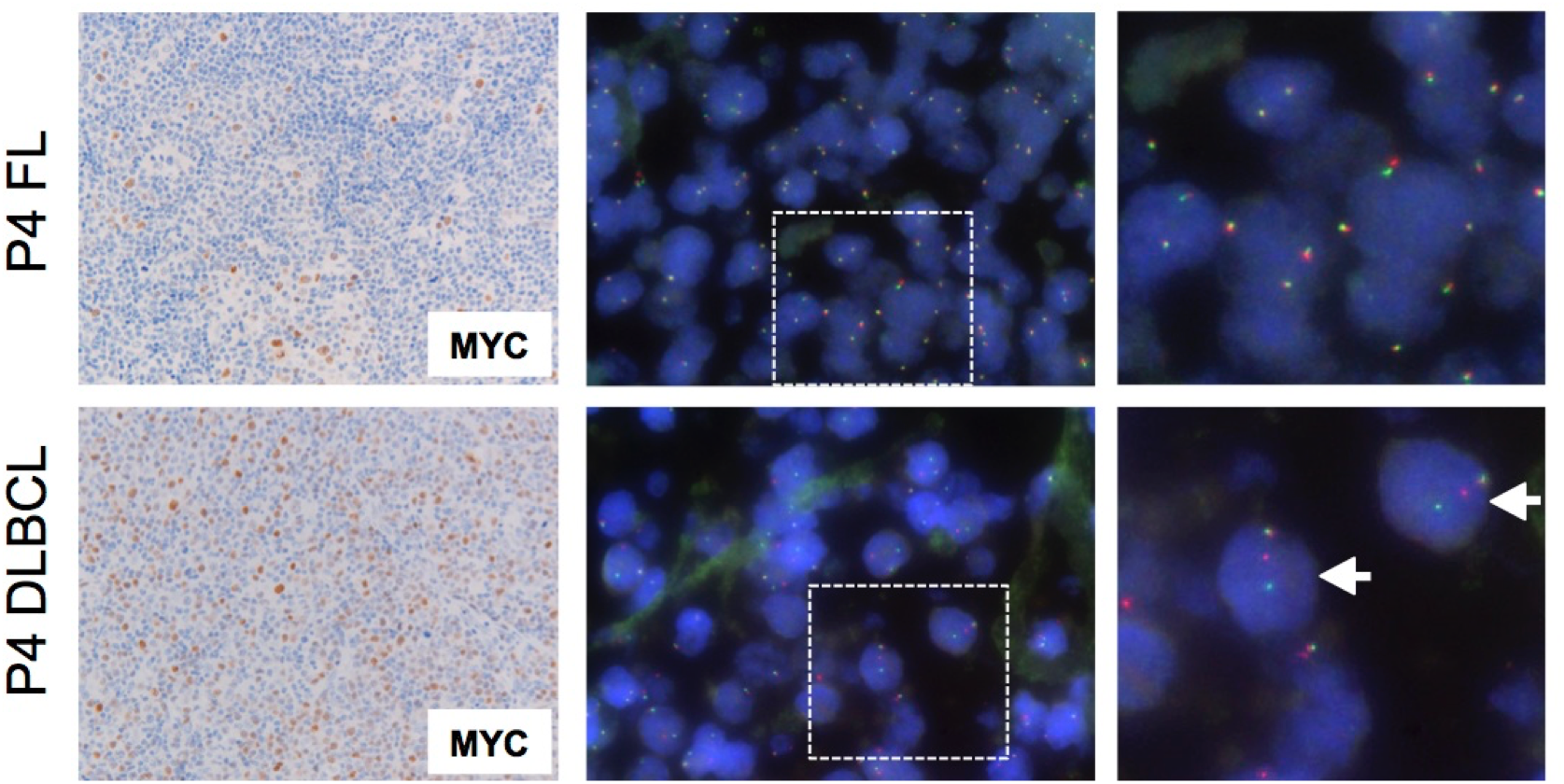
Confirmation of MYC rearrangement in DLBCL. The level of MYC was markedly elevated in the DLBCL of Patient 4 (left panel, at bottom) compared to the FL phase from 2006 (left panel, at top). A break-apart FISH probe revealed rearrangement of *MYC* in the DLBCL of Patient 4, which was not evident in the FL phase (middle panels). A small section of the image (boxed) has been enlarged (shown at right) to highlight the rearrangement (marked with arrows). Representative images are shown.

**Supplementary Figure 4:**
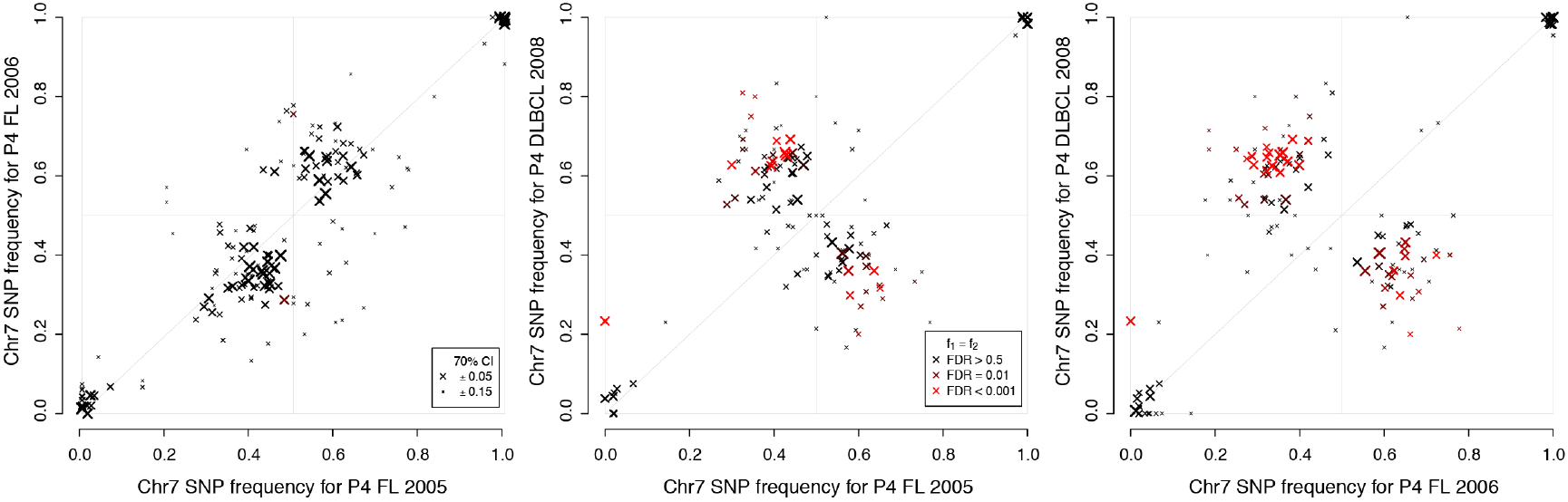
Allele frequencies to track chromosome 7 gain. Allele frequencies are plotted for germline SNPs on chromosome 7 in relapse samples from patient P4. Allele frequencies were compared between each relapse sample. The first panel (at left) shows that P4 FL 2005 and P4 FL 2006 both gained the same chromosome 7. Off-diagonal clusters in the two last panels show that P4 DLBCL 2008 gained the other chromosome 7. The point size and colour represents accuracy and significant difference between the samples, as in Figure 4 (see Supplementary Methods).

**Supplementary Figure 5:**
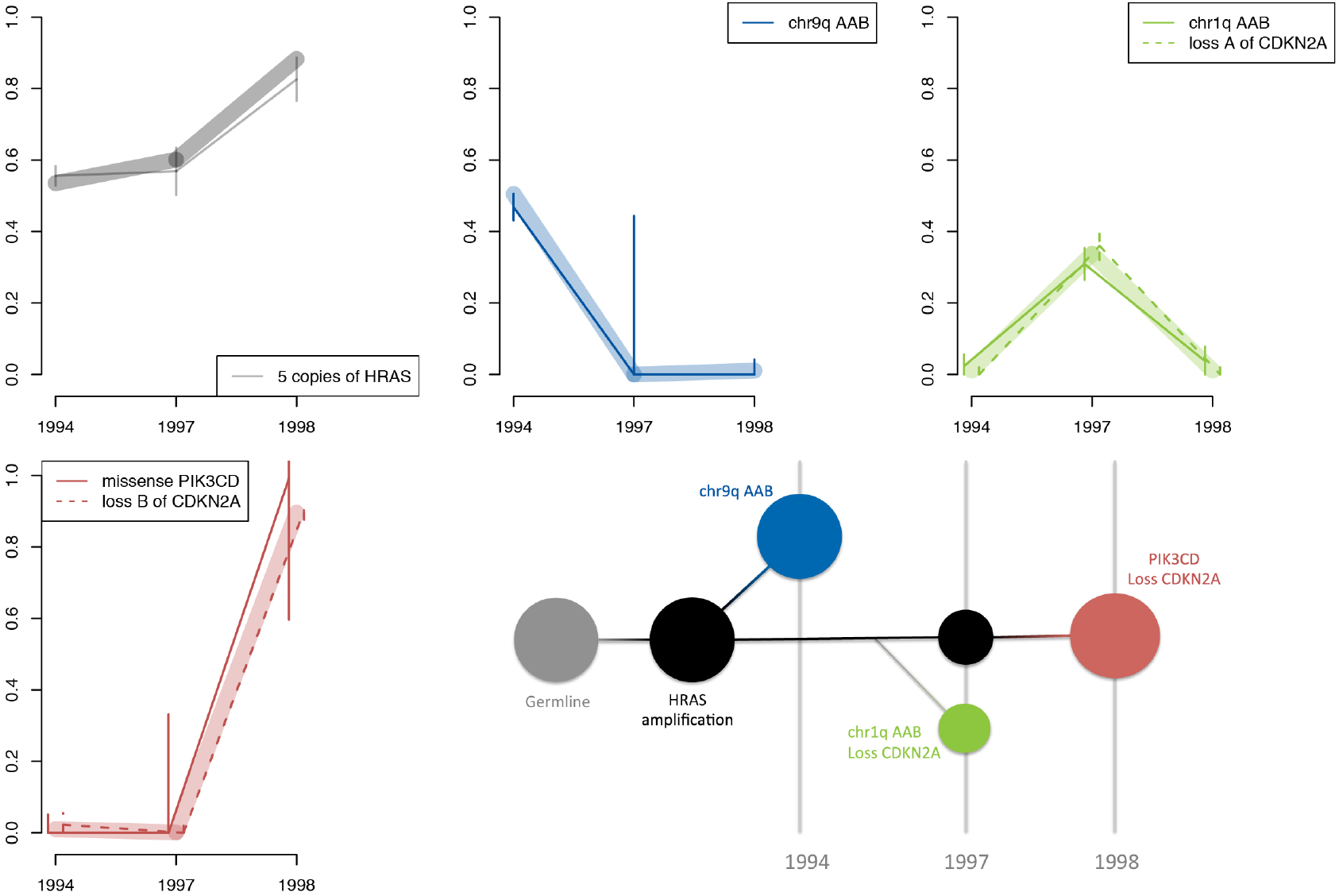
The clonal architecture of samples from P14. Thick lines show the consensus clonalities of a group of mutations, and thin lines highlight individual mutations in that group. Different alleles (denoted as “A” and “B” in the legends) of *CDKN2A* are lost in the 1997 and 1998 samples. The tree shows the evolutionary structure of the cell populations associated with the groups of mutations.

### Integrated analysis of DNA and RNA identifies regulatory mutations that activate key oncogenes

Paired FL and DLBCL samples demonstrated marked differences in gene expression. GSEA was performed to identify cellular processes and pathways altered during transformation **(Supplementary Table 4)**. DLBCL was characterised by high expression of genes associated with cell proliferation (including genes activated during mitosis, E2F targets and MYC targets) **(Supplementary Figure 6)**, in agreement with earlier findings(16, 32).

**Supplementary Figure 6:**
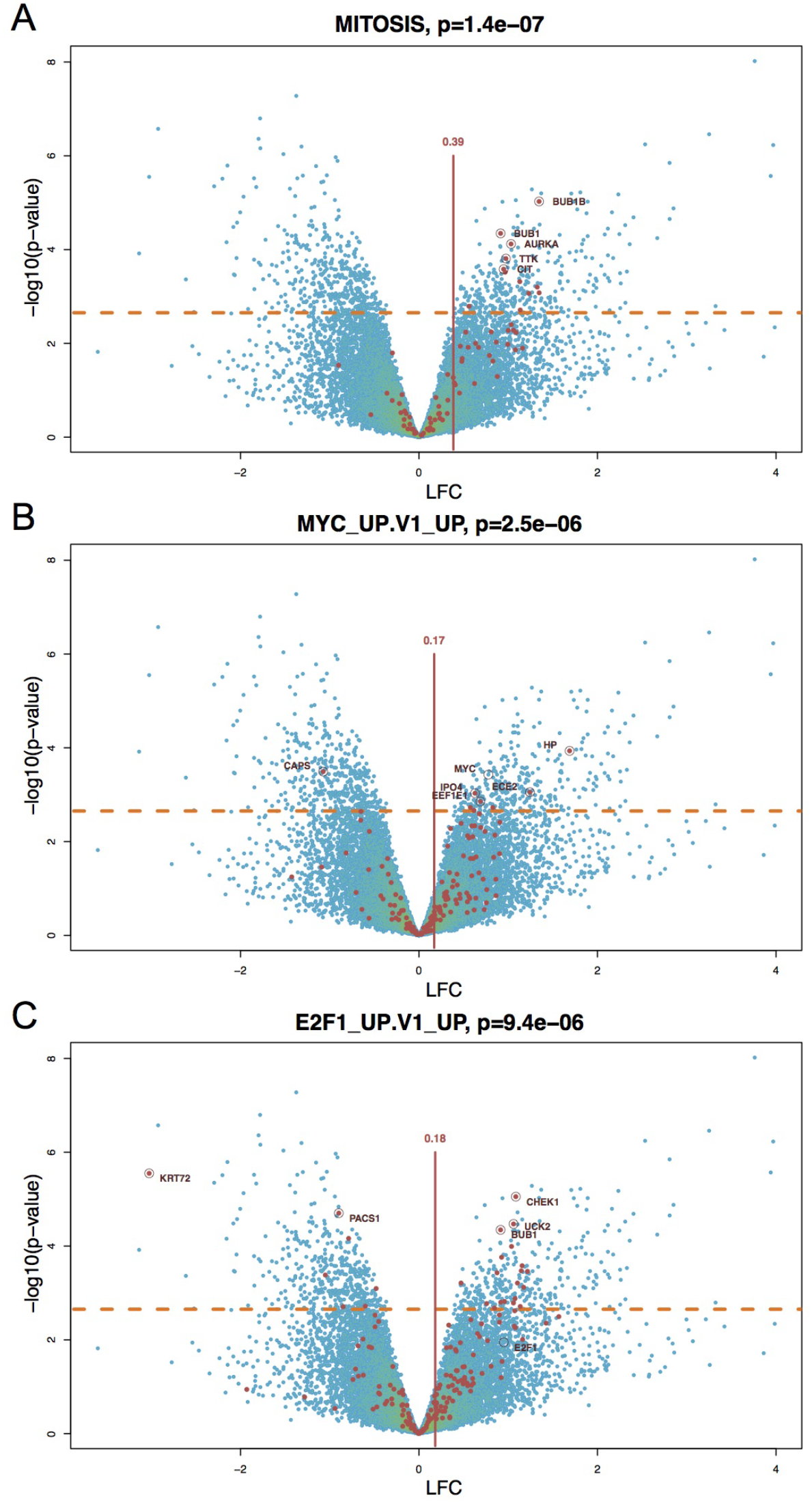
Differential expression between DLBCL and FL. Gene expression was compared between DLBCL and FL. The LFC is plotted for each gene against the statistical significance, with an FDR of 0.05 marked with an orange dotted line. A colour gradient was applied to indicate point density (yellow is dense, blue is sparse). Three illustrative gene sets from the top table in the GSEA are highlighted (A-C). The name of each gene set is displayed, together with the p-value derived from GSEA. The red dots represent the genes in each gene set. The five genes with smallest p-value in each set are circled and labelled. MYC and E2F1 are also highlighted in their respective gene sets. The mean LFC of the genes in each set is shown with a vertical red line.

Integrating DNA and RNA sequencing data allowed us to define mutations that influence transcript abundance and to determine whether they impacted on one, or both, alleles of the gene. We found strong support for predicted SCNVs, including gains in *MYC*, *CCND3* and *HRAS* that were associated with overexpression **(Supplementary Figure 7)**. Loss of *TP53* and *CDKN2A* was also evident at the level of transcript expression.

**Supplementary Figure 7:**
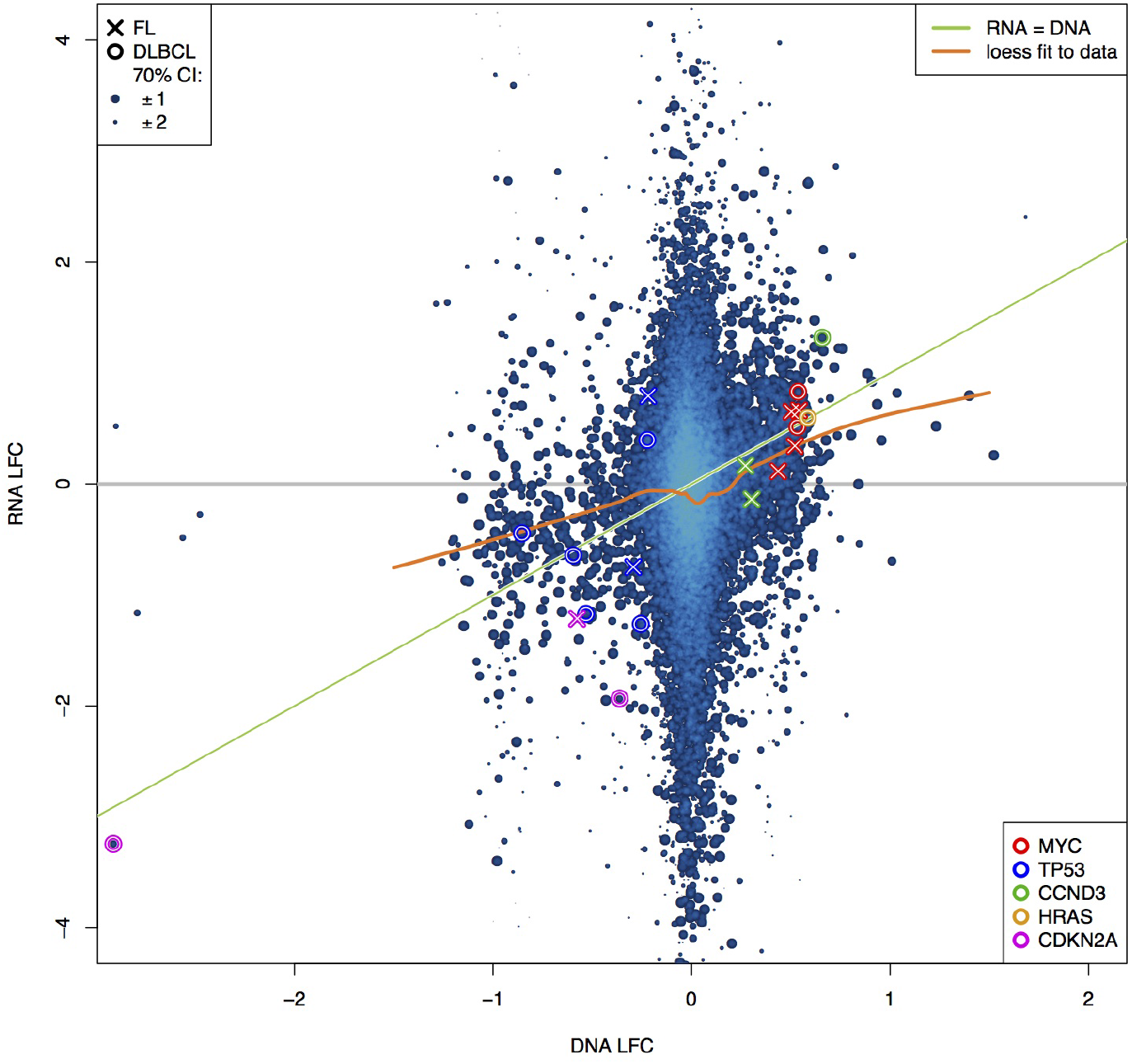
Comparison of DNA coverage with related changes in gene expression. The LFC of the captured genes in the DNA samples compared with the LFC in the RNA. The DNA LFC is taken with respect to the pool of normal samples. The RNA LFC is taken with respect to all other FL samples. Measurements from all patients are presented on the same plot and a colour gradient was applied to indicate point density (yellow is dense, blue is sparse). The point size reflects the accuracy of both measurements (see Supplementary Methods). The green line shows where the RNA LFC is equal to the DNA LFC, representing a complete response of expression on copy number alterations, while the grey line corresponds to expression independent of copy number. The orange line is a weighted loess fit to the data, showing a trend between the green and grey lines, which suggests some buffering of expression levels. Some known cancer genes are highlighted for samples that show signs of altered copy number.

A small number of SNPs were identified that showed significantly different allele frequencies between DNA and RNA. This included a nonsense mutation in *TP53* identified in Patient 6 that was likely targeted by nonsense-mediated decay. Shifted allele ratios in RNA could also be due to mutations that act in cis to drive high-level expression of a single allele, such as mutations or structural rearrangements in promoter elements. Indeed, using this approach we were able to confirm shifted allele frequencies in *MYC* and *BCL2* in the DOHH2 cell line that carries translocations in both genes **(Supplementary Figure 8** and data not shown). Within the patient group we found highly shifted allele frequencies in the RNA of key oncogenes like *MYC* (Patient 4 DLBCL) and *MET* (Patient 7 DLBCL), and in the Rho guanine nucleotide exchange factor (RhoGEF) *TRIO* in two patients (Patient 2 and Patient 8), all of which were accompanied by overexpression **(Figure 4)**. These events were all specific to the DLBCL phase, as they were not present in the matching FL. Oncogenes like *MYC* are up regulated in DLBCL by a range of different mechanisms, including copy number gain, altered microRNA expression and translocations. Immunostaining revealed higher levels of MYC in the DLBCL phase of Patient 4, which could be attributed to a structural rearrangement **(Figure 5)**.

**Supplementary Figure 8:**
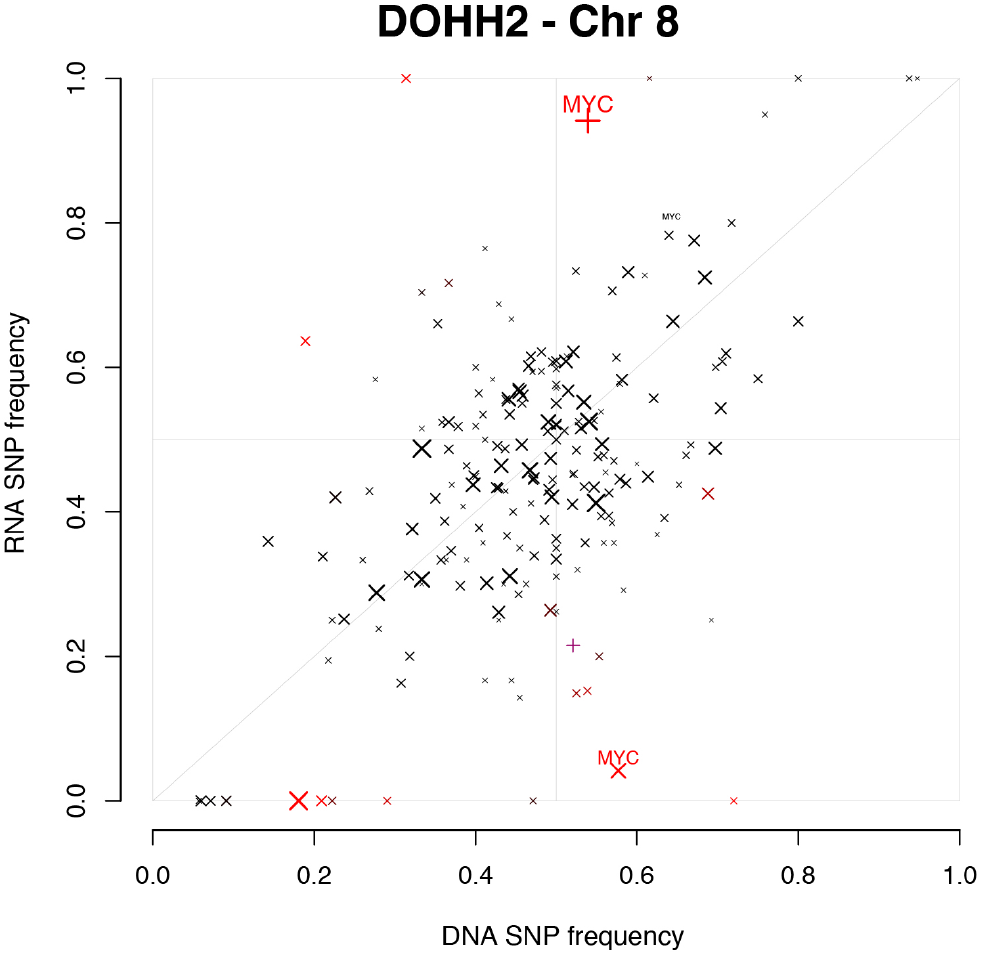
Shifted allele frequencies in RNA for *MYC* in the DOHH2 cell line. The allele frequency is shown for heterozygous germline SNPs in both the DNA and the matched RNA for the DOHH2 cell line. The accuracy of the allele frequency, influenced by coverage, is represented by point size (as in Figure 4). SNPs with significantly different frequencies between DNA and RNA are coloured red.

## Discussion

Transformation from FL to DLBCL is associated with a marked change in cellular phenotype and clinical behaviour. We investigated genomic and transcriptomic - changes in signalling networks that accompany transformation. In agreement with earlier studies, we found that DLBCL typically arises because of new mutations in rare progenitors, rather than direct progression from the dominant FL clone present at diagnosis (13, 14). In addition to mutations in *TP53* and the CDKN2A/CCND pathway, we identified a diverse array of mutations in the PI3K pathway that accompanied transformation. Focusing on individual patients we found evidence of convergent evolution, where mutations in the same gene or pathway occur repeatedly to drive expansion of the cancer.

Our analysis identified frequent mutations in the PI3K pathway in FL and DLBCL, including mutations in *PIK3CD*, *PTEN* and *PIK3R3*. Previously Zhang *et al.* identified mutations in *AKT1*, *MTOR* and *PIK3CD* in DLBCL and went on to demonstrate that mutations in the catalytic domain of PIK3CD stimulate downstream signaling (33). In Patient 4 we identified new mutations in the PI3K pathway associated with each relapse, suggesting that they confer a strong selective advantage. This also suggests that the PI3K pathway is central to disease progression for some patients, and that these individuals may benefit from treatment with inhibitors targeting the pathway. Indeed, DLBCL cell lines that carry activating mutations in MTOR are more sensitive to PI3K pathway inhibitors (33), and clinical trials are currently underway to assess the utility of targeting the PI3K pathway in patients with relapsed, refractory non-Hodgkin’s lymphoma.

By monitoring allele frequencies at heterozygous SNPs we were able to track the genotype of each SCNV, which allowed us to identify genes that were recurrently targeted by mutations in multiple relapse samples from an individual patient. In Patient 14 this identified two distinct events targeting *CDKN2A*. In Patient 4 we demonstrated that chromosome 7 was gained on three separate occasions, and in each case this was coincident with a new mutation in the PI3K pathway. This could suggest that neither event on its own is sufficient to support the outgrowth of the lymphoma clone, and that these events act cooperatively. Further genomic profiling of FL/DLBCL would allow for a more robust assessment of whether there is an interaction between chromosome 7 gain and PI3K pathway mutations.

Integrating both DNA and RNA data allowed us to determine the expression status of each SSNV, revealed nonsense-mediated decay of some transcripts and provided support for predicted SCNVs. This analysis also accurately identified candidate regulatory mutations, even when they occurred outside of capture regions, which were detected based on allele specific changes in expression. These events were all acquired during transformation to DLBCL and were associated with elevated expression of the target gene. We went on to validate the rearrangement of *MYC* in Patient 4 using FISH, and confirmed that the rearrangement was specific to the DLBCL stage. The DLBCL from Patient 4 shares features associated with prior relapses, including mutations in the PI3K pathway and chromosome 7 gain, but the MYC rearrangement appears specific to the transformed disease. Indeed, DLBCLs that carry t(14;18) often carry rearrangements that boost *MYC* expression (34). Allele specific changes were also identified in *MET* and *TRIO*. *MET* was previously found to be up regulated during transformation in paired FL/DLBCL samples (35). Although we identified one patient that acquired a missense SSNV in *MET* upon transformation (Patient 11 DLBCL: Gly326Glu), there is, on the whole, limited evidence for somatic mutations in *MET* in DLBCL. *TRIO* is part of a large family of RhoGEFs. While TRIO has not been implicated directly in DLBCL, there is growing evidence that supports a role for G protein-coupled receptors (such as GNA13 and GNAI2) and RhoA in B cell malignancy (11, 12, 31, 36, 37). Our results warrant further investigation into mechanisms that modulate cell signalling during transformation.

Our analysis has identified a complex pattern of clonal selection during disease progression in FL. In some patients there is clear evidence of convergent evolution within the disease, where the same pathway is targeted time and again, which illuminates core pathways that support the growth of the cancer. This highlights the importance of developing pathway level analyses that accommodate the mutational diversity observed within FL and DLBCL. Ideally these core dependencies could be identified at diagnosis, rather than using multiple relapse samples. Greater understanding of the pathways that drive transformation in FL should enable the development of improved prognostic classifiers and stimulate the development of new, more effective treatments.

## Data Accession

The captured DNA and transcriptome sequencing data, as well as SNV and CNA calls, have been deposited at the European Genome Phenome Archive under accession EGAS00001002175.

## Author Contribution

CF wrote the code and performed the analysis together with input from TS and IJM. CF, IJM and DDJ wrote the manuscript. DDJ, MH and REK contributed and assessed clinical samples, and AB, DDJ, RJCK and IJM performed experiments. All authors read and approved the final manuscript.

## Acknowledgements

We wish to thank the members Genomics Facility and the Molecular Pathology & Biobanking Service at the NKI. Bauke YIstra provided valuable feedback on the methods component. This work was made possible through Victorian State Government Operational Infrastructure Support and Australian Government NHMRC IRIISS (9000220) and by an NHMRC Program Grant (1016647) and Fellowships (575581 to IJM, 1058344 to WSA). We also wish to acknowledge the generous support of Mr. Malcolm Broomhead who provided philanthropic support for the research.

## Supplementary Data

### Supplementary Tables

**Supplementary table 1: Clinical features of the patient cohort, including interval between diagnosis of FL and DLBCL.** Treatment abbreviations: CHLOR – Chlorambucil, CHOP – Cyclophosphamide, Doxorubicin, Vincristine, Prednisone, CHP – Cyclophosphamide, Doxorubicin, Prednisone, CVP – Cyclophosphamide, Vincristine and Prednisolone, DHAP – Dexamethasone, Cytarabine, Cisplatin. ETOP – Etoposide, FLUDA – Fludarabine, IFOS – Ifosfamide, PROMACE – Prednisone, Methotrexate, Doxorubicin, Cyclophosphamide, and Epipodophyllotoxin, R – Rituximab, RT – Radiation Therapy, VIM – Etoposide, Ifosfamide and Mitoxantrone, W&S – Wait and see (monitor). Status: AWD – Alive with disease, CR – Complete remission, DOD – Died of disease, DOOC – died of other causes.

**Supplementary Table 2: Capture set and expression status.** The genes in the capture set are listed in the first column, with the chromosome, start and end positions of the gene in hg19 coordinates in column 2 to 4. Column 5 lists the mean expression over all samples with RNA sequencing data in reads per million per kbp (RPKM). Columns 6 to 31 show gene expression values for each individual sample in RPKM. Column 32 to 57 show the uncorrected read counts from featureCounts, including only reads with a mapping quality of 10 or more.

**Supplementary table 3: Mutation description across FL and DLBCL cohort**. This table contains all SSNVs and SCNVs contributing to **Figure 1**.

### For SSNVs the columns are

- **Patient:** which patient the mutation is detected in.
- **Chr, start, end, gene:** The genomic location in hg19 coordinates and the gene in which the mutation is located. For indels, this refers to the base immediately before the mutation.
- **Reference, variant:** The reference and variant bases.
- **Type, exon, AApos, AAbefore, AAafter, domain, polyPhen, sift:** output from the variant effect predictor (VEP), showing the variant type, which exon the variant is located on, the amino acid position, reference and variant, the domain the variant is located in, and estimates of the functional impact of the mutation by polyPhen and SIFT.
- **FL count, FL depth, FL clonality, FL clonality uncertainty:** Prevalence of the mutation in the diagnosis FL sample. The columns are the number of reads supporting the variant, the total read depth over the position, the estimates clonality (using local copy number information) and 70% confidence interval.
- **DLBCL count, DLBCL depth, DLBCL clonality, DLBCL clonality uncertainty:** same as above for the DLBCL sample of the patient.
- **Classification:** How the mutation is classified in Figure 1: shared, FL specific or DLBCL specific. SCNVs are described by:
- **Patient:** which patient the mutation is detected in.
- **Chr, start, end:** The genomic region in hg19 coordinates that have evidence of the SCNV. Note that there are typically large gaps between the captured genes, which limits the accuracy of the break point estimate. The coordinates provided reflect the last capture regions that show evidence of the SCNV.
- **FL call, FL clonality, FL clonality uncertainty:** The called SCNV by superFreq in the FL sample. “A” and “B” represent the paternal and maternal alleles, so that for example “AB” is normal diploid, “AAB” is gain, “AA” is CNN-LOH, “A” is loss of one allele. The estimated clonality and 70% confidence interval are calculated from the coverage LFC and SNP frequencies in the region.
- **DLBCL call, DLBCL clonality, DLBCL clonality uncertainty:** as above for the DLBCL sample of the patient.
- **Classification:** How the mutation is classified in Figure 1: shared, FL specific or DLBCL specific.

## Supplementary Methods

### Point size by accuracy

In a number of figures **(Figure 4 and Supplementary Figure 1, 4 & 8)** we have scaled point size to reflect the accuracy of the measurement, which reflects the number of reads the measurement is based on.

In the middle and lower panels of Figure 4, the area of each cross (the square of the linear size), is proportional to the geometric mean of the coverage over the SNP in the two samples that are being compared. An upper limit is set at a mean read depth of 225, over which the point size is not further increased.

In Supplementary Figure 1, the point size of the coverage LFC (top panels) and allele frequency (bottom panels) are determined by the confidence interval calculated by superFreq, which is also shown with error bars. The confidence interval of the coverage LFC is calculated using limma-voom, with an added penalty if the sample shows larger fluctuations between neighbouring genes than is expected from the limma-voom estimates. The confidence interval of the allele frequency is based on the summed coverage of the heterozygous germline SNPs in the region, or on the region where the log likelihood (calculated with a binomial distribution for each SNP, combined with Fisher’s method) is within 1% of maximum. The larger of the two confidence intervals is used.

In Supplementary Figure 7, the confidence interval in the DNA LFC is taken from superFreq as in Supplementary Figure 1, while the confidence interval of RNA LFC is estimated using limma-voom. The ranges of the DNA and RNA confidence intervals are added in square for each point and used to determine point size. The linear sizes of the points are inversely proportional to the joint confidence interval, with a maximum size at a joint confidence interval of 0.5. These sizes are also used as weights in the loess fit of the data. Note that most points have a larger variance in the RNA LFC, which dominates the uncertainty estimate.

The confidence intervals are set to 70% to correspond to the standard deviation of a normal distribution. This does not imply that the error profiles of the measurements are Gaussian.

### Settings for alignment and variant calling

- DNA reads were aligned to hg19 using bwa mem v0.7.10-r789 mem with default settings.
- RNA reads were aligned with Tophat v2.0.12 with options ––read-realign-edit-dist 0 and ––b2-very-sensitive.
- Variant detection was performed with samtools version 0.1.19-44428cd with settings - q 1 - Q 15 - A, followed by Varscan v2.3.6 mpileup2cns with settings ––strand-filter 0 ––p-value 0.01 ––min-var-freq 0.01. This preliminary set of variants was filtered using superFreq.
- SSNVs were assessed by Ensembl Variant Effect Predictor version 75 (http://www.ensembl.org/info/docs/tools/vep) with the ––everything option.

### superFreq

We developed a new analysis pipeline called superFreq that is specifically designed to track clonal relationships in cancer. The method is suitable for capture datasets, including whole exomes. The latest version of superFreq can be found on Github:

https://github.com/ChristofferFlensburg/superFreq.

Some of the main issues we wanted to address with this method were
1. We wanted to be able to generate SCNV calls from coverage and allele frequencies without using a matched normal.
2. We required a low false positive rate for SCNV calls, but with sensitivity for deletions and amplifications of individual genes.
3. We needed a method that could track allele specific SCNVs over multiple samples.
4. We wanted to track the quality of both the SSNV and SCNV calls and use this information to inform clonality estimates.

To achieve these goals, we used the following approaches
1. In the presence of a matched normal, the heterozygous SNPs are called from the normal sample. When a matched normal was not available we identify heterozygous SNPs from the cancer sample directly.
2. Read coverage is compared to reference normal samples from the same platform. This removes the requirement of a matched normal, and allows us to estimate biological and technical variance between samples.
3. To decrease the rate of false calls, it is essential to account for as many systematic error sources as possible and to empirically assess the variance estimates. We perform standard GC correction and MA normalisation of the read counts over the exons, as well as a correction of the sex chromosome counts. We merged read counts by gene to reduce the variance and limit the number of false positive calls. Coverage LFCs are used together with SNP frequencies and their uncertainties to cluster regions and derive allele specific copy numbers.
4. Tracking the genotype of each SCNV allows us to identify instances where genes were targeted by independent events in multiple samples from an individual patient.
5. Our approach incorporates the level of uncertainty calculated for each mutation when constructing clonal relationships. The most reliable mutations, i.e. those with the lowest variance (typically high coverage SSNVs in diploid region and SCNVs over large genomic regions), are used to anchor the clustering.

A more detailed technical evaluation of superFreq is in preparation. The version of superFreq used in this publication is available: https://github.com/ChristofferFlensburg/cnv-caller/commit/677cf2e59ccefd18d0ce71833045df92e35905df. The supplementary file generateFigures.R was used to generate the figures in this publication from the superFreq output and has been deposited with the data at EGA.

### Three independent chromosome 7 gains in Patient 4

Chromosome 7 is gained in the last three time points of Patient 4. By assessing the genotype of the SCNV (tracking the chromosome that is gained) and considering the other mutations present, we established that the gain of chromosome 7 occurred on three separate occasions.

To show this, we first observe that the 2008 DLBCL sample has gained a different allele of chromosome 7 from the other samples, which is evident in the SNP allele frequency scatter plots **(Supplementary Figure 4)**. This shows that the gain of chromosome 7 in 2008 DLBCL is a separate event from the 2005 FL and 2006 FL. The 2005 FL and 2006 FL gained the same allele, but this cannot originate from the same mutation, as that would be inconsistent with the behaviour of many other mutations. Clone A and Clone C are both subclones of Clone B, but we know that the classification of one of the clones is incorrect because in the 2005 FL the sum of the clonality measures for Clone A (~0.35) and Clone C (~0.35) exceeds the clonality of Clone B (~0.5) **(SM Figure 1, below)**. Clone A cannot be a subclone of Clone C, as the 2006 FL shows Clone A, but not Clone C. Similarly, Clone C cannot be a subclone of Clone A, as the 2008 DLBCL shows Clone C, but not Clone A. This is a nonsensical result, as any two cell populations must be either disjoint, or one must be a subclone of the other.

To resolve this issue we must break apart one of the groups - Clone A, Clone B, or Clone C. Splitting up Clone C would imply that the four SSNVs and the CNN-LOH of 11q occurred twice independently, which is extremely unlikely **(SM Figure 2, below)**. Similarly, Clone B cannot be split up, as it is supported by three SSNVs **(SM Figure 3, below)**. We are left with the only plausible explanation that Clone A is actually two clones, Clone A1 present in the 2005 FL only, and Clone A2 present in the 2006 FL only **(SM Figure 4 and 5, below)**. The gain of chromosome 7 in clone A1 and A2 was originally combined because they gained the same allele of chromosome 7. Splitting Clone A requires that a single event, the gain of chromosome 7, occurred twice, rather than a host of SSNVs. No SSNVs were identified that showed behaviour similar to the original Clone A (present in 2005 FL and 2006 FL, but not in 2008 DLBCL), while there are numerous mutations supporting clone A1 and A2, which further supports our interpretation of the clonal architecture **(Figure 3 in the manuscript)**.

**SM Figure 1:**
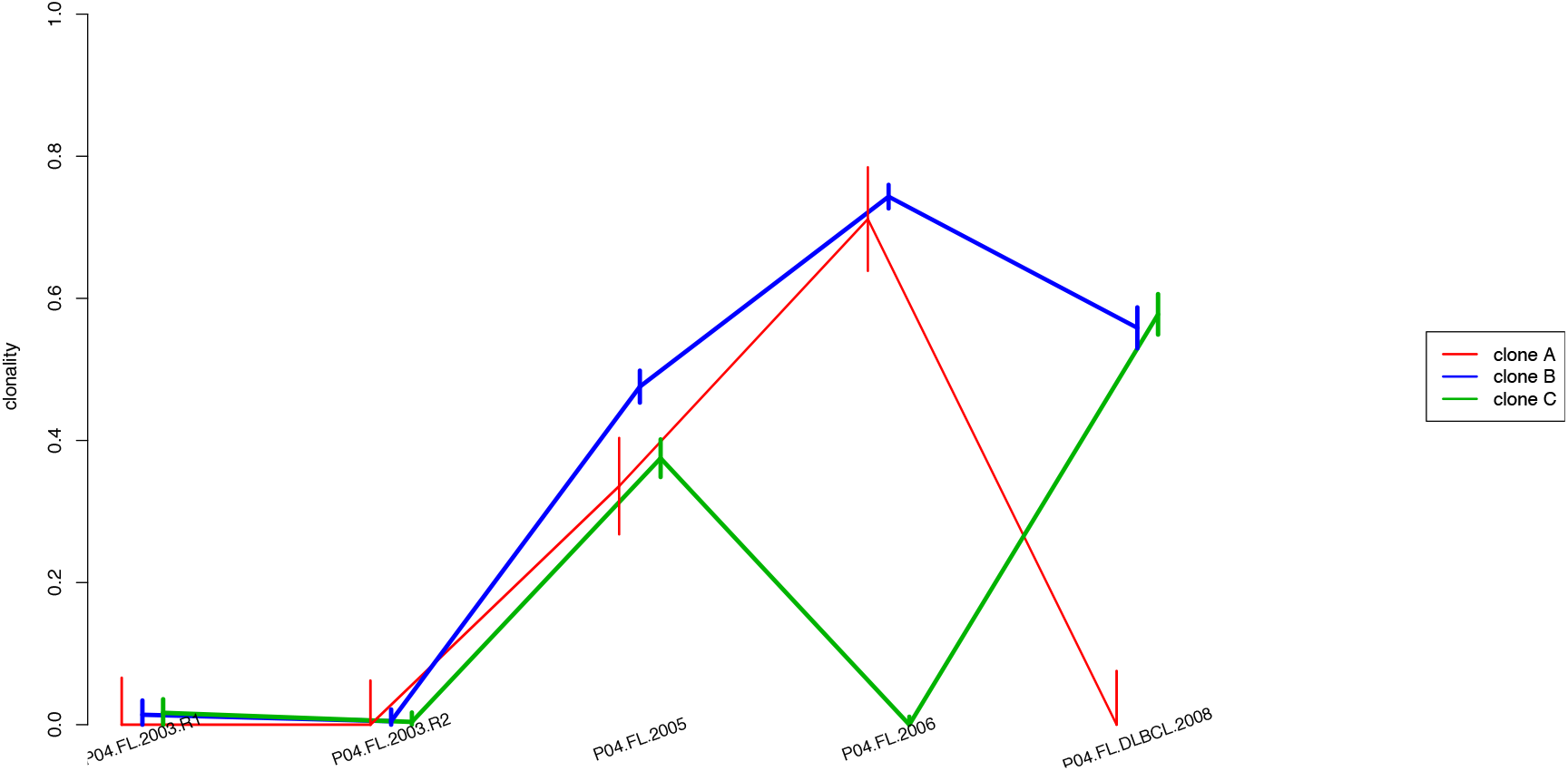
clone A, B and C as called by superFreq.

**SM Figure 2:**
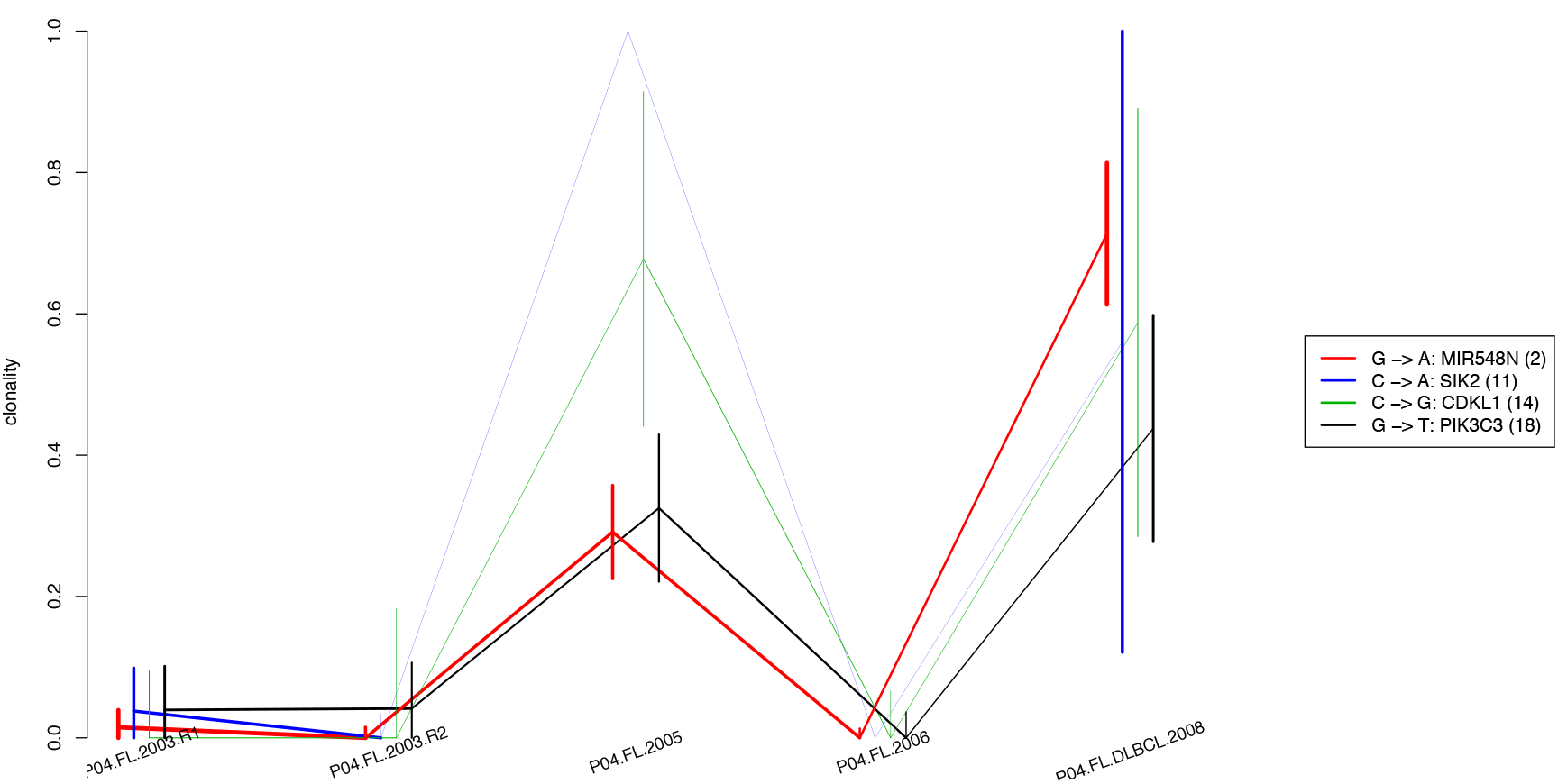
Supporting SSNVs for clone C.

**SM Figure 3:**
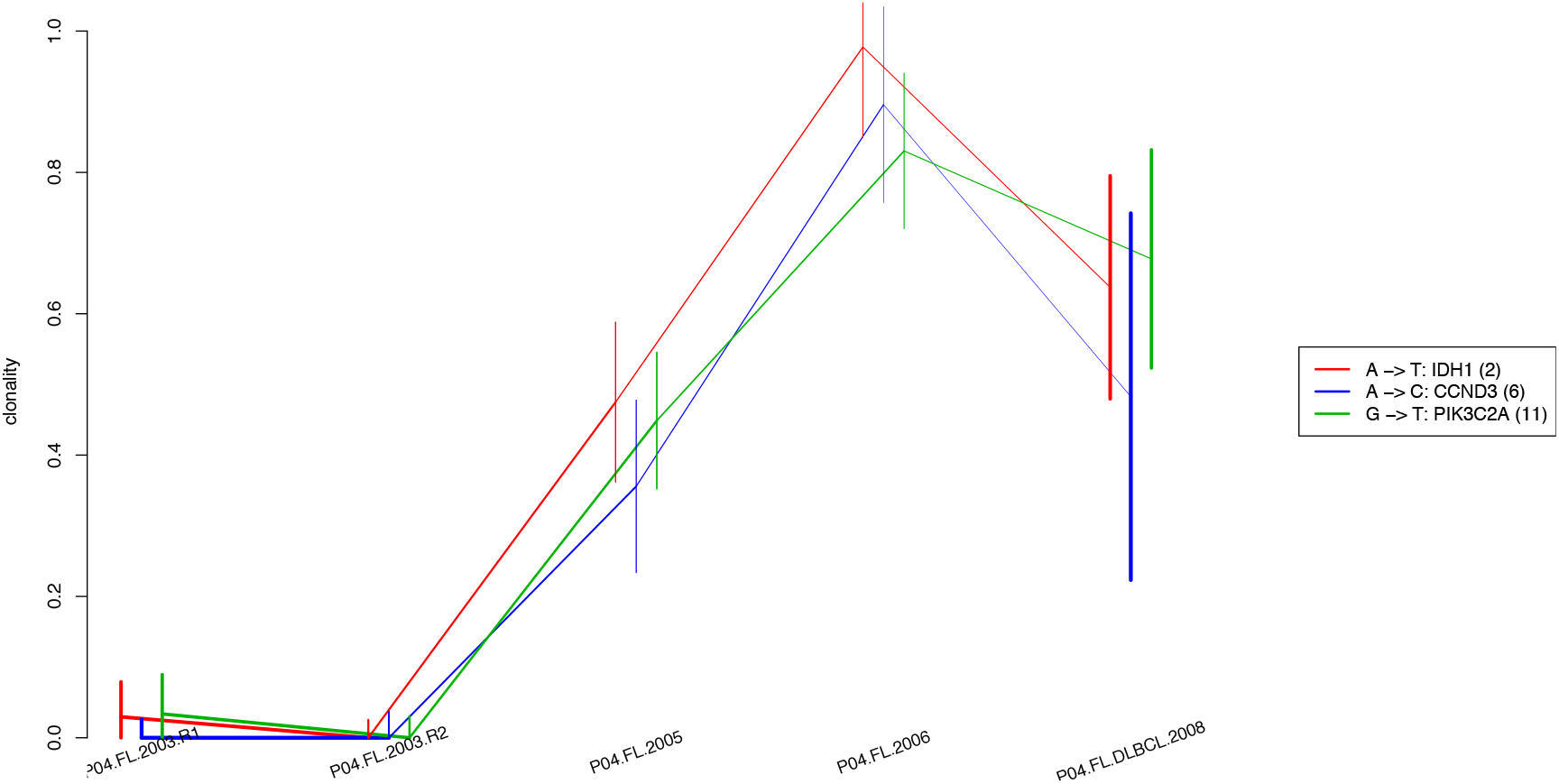
Supporting SSNVs for clone B.

**SM Figure 4:**
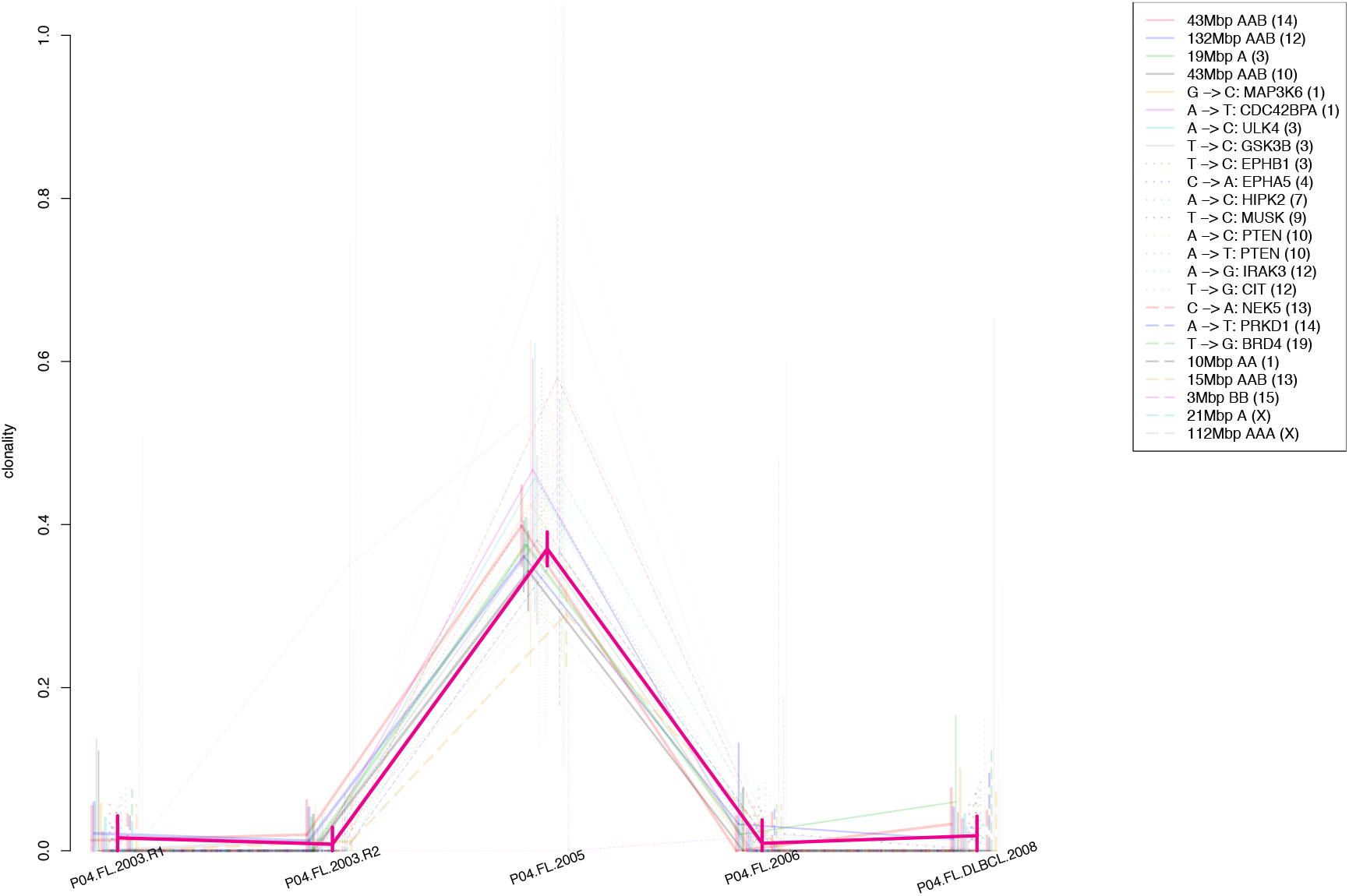
Supporting mutations for clone A1.

**SM Figure 5:**
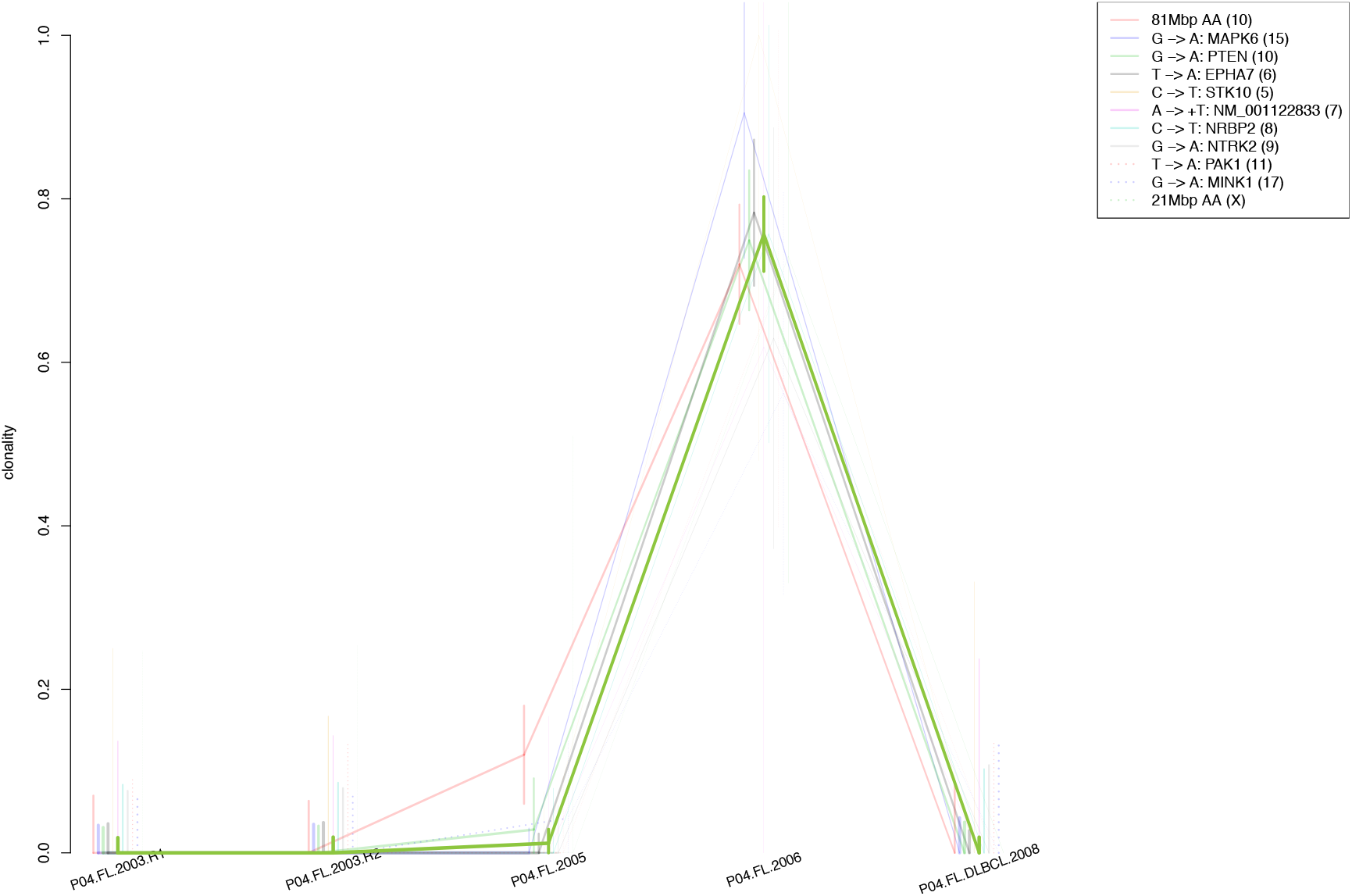
Supporting mutations for clone A2.

